# MaSS-Simulator: A highly configurable MS/MS simulator for generating test datasets for big data algorithms

**DOI:** 10.1101/302489

**Authors:** Muaaz Gul Awan, Fahad Saeed

## Abstract

Mass Spectrometry (MS) based proteomics has become an essential tool in the study of proteins. The big data from MS machines has led to the development of novel serial and parallel algorithmic tools. However, the absence of data benchmarks and ground truth makes the algorithmic integrity testing and reproducibility a challenging problem. To this end, we present MaSS-Simulator, which is an easy to use simulator and can be configured to generate MS/MS datasets for a wide variety of conditions with known ground truths. MaSS-Simulator offers a large number of configuration options to simulate control datasets with desired properties thus enabling rigorous and large scale algorithmic testing. We assessed 8,031 spectra generated by MaSS-Simulator by comparing them against the experimentally generated spectra of same peptides. Our results showed that MaSS-Simulator generated spectra were very close to the real-experimental spectra and had a relative-error distribution centered around 25%. In contrast the theoretical spectra for same peptides had relative-error distribution centered around 150%. Source code, executables and a user manual can be downloaded from https://github.com/pcdslab/MaSS-Simulator

## 1 Introduction

High performance liquid chromatography (HPLC) combined with tandem mass spectrometry has revolutionized the study of proteins. It has become an essential part of systems biology studies ([1]), drug discovery research ([2]), detection and determination of phenotypes of cancer ([3]), toxicology studies ([4]) and evolutionary biology ([5]). A usual mass spectrometry (MS) based proteomics pipeline consists of breakdown of unknown proteins into smaller chains known as peptides and proceeds by separating them using high-performance liquid chromatography (HPLC). Separated peptides are then transferred to a Mass Spectrometer to obtain MS1 spectra ([6]). In the fragmentation process each unknown peptide is broken down into several types of ions to yield an MS/MS spectrum. Each ion in MS/MS spectrum is represented by its mass-to-charge ratio and corresponding intensity to represent its relative abundance. Collision Induced Dissociation (CID), Higher Energy Collisional Dissociation (HCD) ([7]), Electron Capture Dissociation (ECD), and Electron Transfer Dissociation (ETD) are common dissociation strategies ([8]) ([6]). The data generated by the Mass Spectrometers is processed using a software pipeline([9]). This pipeline relies heavily on the accuracy of the peptide deduction algorithms. These algorithms ([10]) were either designed for MS/MS spectra generated by a specific ion-dissociation strategy or have only been tested on very limited sets of data. Comparing and assessing the performance of these large number of algorithms is a challenging problem due to the lack of systematic data generation where the parameters of benchmarks are in control of the method developer ([11]). These parameters may include noise content, quality of spectra, different peptide coverage, and varying dissociation strategies.

Generating experimental spectra is a costly process with many parameters not in one’s control. One way of obtaining MS data sets in which all the parameters are in control of the method developer is with the help of simulators. Such simulators have been used successfully for generation of next generation sequencing data ([12]). Existing simulators for MS data, such as MSSimulator ([11]) can be used to simulate LC-MS experiments but this simulator offers a very limited control over the simulation of MS/MS spectra. Similarly, MS-Simulator ([13]) can generate theoretical spectra with accurate y-ion intensities for a handful of experimental conditions. These simulators either offer very limited control over the parameters or use pre-trained models specific for a particular instrument or ion-dissociation strategy to simulate MS/MS spectra.

To the best of our knowledge there does not exist a simulator for MS/MS data which will allow careful exploration of the space of the parameters associated with MS/MS data. Such exploration will allow one to identify bottlenecks, strengths and weaknesses in the proposed algorithms for MS based proteomics. In this paper we introduce MaSS-Simulator, which offers many configurable options for generating controlled MS/MS spectra. By correctly configuring this simulator with simple configuration file control datasets with desired properties and ground truth peptides can be obtained and used for assessment of proteomics algorithms.

## 2 Implementation

Here we discuss the implementation and features offered by MaSS-Simulator.

### 2.1 Controlled Generation of Ions

Fragmentation process which leads to the generation of MS2 spectra is highly dependent upon the ion-dissociation technique, instrument and other factors ([8]). To give user a complete control over the conditions of fragmentation we introduce a feature of *Ion Generation Probability* (IGP). IGP value for each ion determines the likelihood that a given ion will be generated in the simulation. Using the ion generation probabilities peptide coverage can be controlled. Hence by correctly selecting the ion series and their corresponding IGP values, any dissociation strategy can be simulated. Immonium ions may be formed for some ion dissociation techniques which are helpful in detecting certain amino acids ([10]). MaSS-Simulator can be configured to generate these ions with a given IGP value.

### 2.2 Controlling the Relative Abundance Values

A lot of effort has been made to predict the ion intensities theoretically but the developed models have been trained only for a handful of experimental conditions ([13]). For our case we used average of relative intensity values as default settings e.g. average intensity of y ions is usually two times that of b ions ([14]). Intensity values for each ion series can also be adjusted by the user from the configuration file.

### 2.3 Static and Variable Post Translational Modifications

MaSS-Simulator is capable of simulating both the static and variable Post Translational Modifications (PTM). All desired types of modifications can be listed in the *modifications.ptm* file and the details on the format of the file are given in the user manual. For our experiments we tested both static Carbamidomethyl: C+57.021) and variable modifications (Phosphorylation: STY + 79.966, Deamidation: NQ+0.984 and Oxidation: M+15.995).

### 2.4 Sound to Noise Ratio

In most MS2 spectra about 90% of the data is noise/non-essential peaks ([9]). Nature and amount of noise in spectra can vary greatly with the experimental conditions. MaSS-Simulator gives an option to add *random* noise peaks in spectra that can either be *uniformly* distributed or follow a *Gaussian* distribution with the possibility of including a user defined noise model. To control the amount of noise to in spectra, we use SNR (Sound to Noise Ratio) value given by:

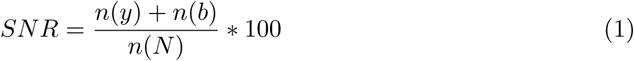

Where *n*(*y*) and *n*(*b*) are the number of y and b ions respectively, while *n*(*N*) is the number of noise peaks.

### 2.5 File Output format

The simulated spectra are output in MS2 format which can be conveniently converted to any desired format using proteowizard ([15]).

## 3 Experimental Datasets

To perform the assessment of simulated spectra generated by MaSS simulator we made use of experimentally generated spectra from 8,031 peptides obtained using two different ion-dissociation strategies.

### 3.1 Control Dataset Generation

To perform a rigorous assessment of MaSS simulator we needed datasets with varying ion-dissociation strategies. To do so we opted for the same datasets we used for the assessment of MS-REDUCE algorithm [9]. To obtain these datasets we minced and sonicated a freshly isolated rat liver in guanidine-HCL(6M, 3ml). To 500g aliquots of liver sample (before trypsinization), a peptide standard corresponding to C-terminal sequence of the water channel Aquaporin-2 from rat, (Biotin-LCCEPDTDWEEREVRRRQS^∗^VELHS^∗^PQSLPRGSKA) phosphorylated at S256 and S261 were added with distinct amounts of 0.2 nmol, 20 pmol and 2 pmol. We will further refer these samples as DS-1, DS-2 and DS-3 respectively.

Same procedure was then repeated for AQP2 peptide standard (Biotin-LCCEPDTDWEERE-VRRRQSVELHSPQS^∗^LPRGSKA) phosphorylated at S264, with 0.2nmol, 20pmol and 2pmol amounts. The same procedure as above was repeated for another AQP2 peptide standard (Biotin-LCCEPDTDWEEREVRRRQSVELHSPQS^∗^LPRGSKA) phosphorylated at S264, with amounts of 0.2 nmol, 20 pmol and 2 pmol. These samples will be referred to as DS-4, DS-5 and DS-6 respectively in future. Peptides samples were then desalted and suspended in 0.1 percent formic acid before being analyzed by a mass spectrometer. We used HCD and CID fragmentation strategies on each of the above datasets thus yielding a total of 12 datasets. Throughout this paper each dataset can be identified by the fragmentation strategy followed by the name of peptide sample i.e. HCD-DS-5 means the mass spectra dataset obtained after performing HCD fragmentation on DS-5 peptide sample.

All the above datasets were then searched against a rat proteome database obtained from (http://uniprot.org) using the Tide software [16]. To perform this search a static modification of carbamidomethyl (C:+57.021 Da) and dynamic modifications of Phosphorylation (S,T,Y: +79.966), Deamidation (N,Q: +0.984) and Oxidation (M: +15.995) is included. The resulting Peptide Spectral Matches (PSMs) are processed through decoy database based FDR filtering method using the software percolator [17]. Only the PSMs having an FDR value of less than 0.01 are kept and remaining are discarded. Using this filtering mechanism we obtained a total of 4,076 spectra with CID fragmentation with their corresponding peptides and 3,995 spectra with HCD fragmentation with their corresponding peptides. In total we obtained 8,031 peptides for which we had the corresponding experimental spectra. We call these PSMs as control peptides and control spectra.

### 3.2 Analysis of Experimental Spectra

The above obtained control peptides along with their experimental spectra were analyzed to determine the peptide coverage and SNR value. To compute the peptide coverage, we used the following equation:

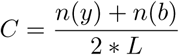

here n(b) and n(y) represent the number of b and y ions in the spectrum while *L* is the length of the corresponding peptide. Similarly, we computed the SNR value for each spectrum using the following equation:

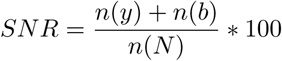

Where *n*(*y*) is the number of y-ions, *n*(*b*) is the number of b-ions while *n*(*N*) is the number of noise peaks to be added.

Now our control dataset consists of a peptide, a corresponding experimental spectrum, its peptide coverage and SNR values. The peptide coverage and SNR values will be used to configure the MaSS simulator when simulating spectra for each peptide.

## 4 Quality Assessment of Simulated Spectra

To assess the spectra generated by MaSS Simulator, we shortlisted 8,031 control peptides for which we had high confidence experimental spectra available. Detailed process of generation of experimental data has been discussed in Section 3 and the work-flow for shortlisting the experimental spectra has been visualized in Fig. 1 (A). The list of control peptides along with a configuration file containing the SNR and coverage values and a *modification.ptm* was given as input to the simulator as shown in Fig. 1 (B). In the configuration file we used peptide coverage values as the IGP value for b and y ion series and the SNR values to control the amount of noise. Other parameters in configuration file for this experiment can be found in the Table 1. At the output we obtained simulated spectra for each control peptide.

**Figure 1:**
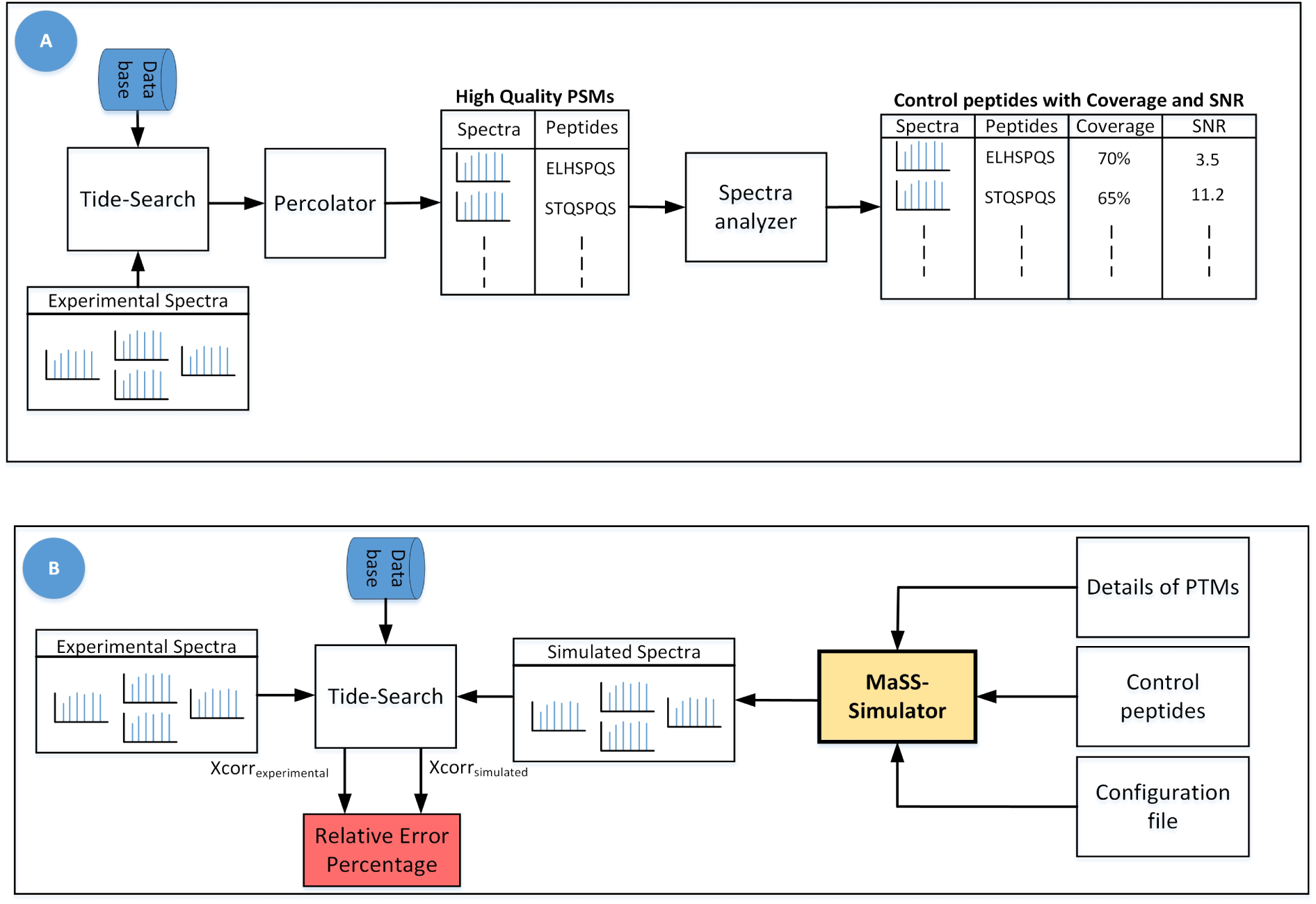
Figure A) shows the workflow used for obtaining experimental spectra with high confidence PSMs along with their Coverage and SNR values. Figure B) shows the workflow for generation and assessment of simulated spectra. To determine the relative error percentage for theoretical spectra, we replaced simulated spectra with theoretical spectra in this workflow.

**Table 1:** Table showing default parameters used in configuration file. Values in brackets followed by each ion shows the IGP value. BOD stands for the value based on the data from the analysis of experimental spectra. Since in our study each experimental spectrum had different IGP values, we had to use different value for each spectrum. It can be observed that we have kept the tendency of CID dissociation having a-ions half of that of HCD. In the simulation all the intensity values are multiplied by a factor of 1000

| Parameters for Configuration File |  |  |
| --- | --- | --- |
| Ions(HCD) | Ions(CID) | Intensities |
| b(BOD) | b(BOD) | 1 |
| b-NH<sub>3</sub>(BOD) | b-NH<sub>3</sub>(BOD) | 0.5 |
| b-H<sub>2</sub>O(BOD) | b-H<sub>2</sub>O(BOD) | 0.5 |
| y(BOD) | y(BOD) | 1.75 |
| y-NH<sub>3</sub>(BOD) | y-NH<sub>3</sub>(BOD) | 0.5 |
| y-H<sub>2</sub>O(BOD) | y-H<sub>2</sub>O(BOD) | 0.5 |
| a(BOD) | a(BOD/2) | 0.5 |
| a-NH<sub>3</sub>(BOD) | a-NH<sub>3</sub>(BOD/2) | 0.25 |
| a-H<sub>2</sub>O(BOD) | a-H<sub>2</sub>O(BOD/2) | 0.25 |
| Immonium Ions<br>(75) | Immonium Ions<br>(40) | 0.5 |

Ideally the simulated spectra should closely match the experimentally generated spectra. To assess the similarity between the two sets of spectra we use the work-flow given in Fig. 1 (B). The idea is to compare both the experimental and simulated spectra using a proven method/score. The xcorr scores obtained from Tide database search software [16] give a measure of how closely the spectrum under consideration matches the theoretical spectrum of a particular peptide. We consider the xcorr value for a given experimental spectrum *A* obtained from Tide (when the target peptide is correct) to be a gold standard. For any spectrum to be similar to *A*, it has to attain a similar Tide xcorr score while giving the same target peptide. hence the difference between the two xcorr scores can give us a measure of difference between two spectra. Using the following equation we can compute a relative error percentage, a smaller error means two spectra match closely.

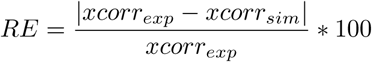

here *xcorr_exp_* represents the xcorr score for experimental spectra and *xcorr_sim_* represents the xcorr score for simulated spectra.

Following the above discussed method, the simulated and the experimental spectra are first searched against the rat database using Tide [16] algorithm which outputs the list of peptide spectral matches (PSMs) and xcorr scores for each set of input spectra. We consider the PSMs from both sets which have the same target peptide and use their xcorr values to compute a relative error percentage using the above equation. The same procedure is repeated by replacing the simulated spectra with simple theoretical spectra and relative error percentage is computed using the above equation by replacing *xcorr_sim_* with *xcorr_theo_* which represents the xcorr score for theoretical spectra.

A boxplot is used to compare the relative error distributions for simulated and theoretical spectra as shown in Fig. 2 through 5. To reduce the number of plots we grouped together all the experimental datasets in four groups. Two groups each for each ion-dissociation strategy, one group only had those spectra which were obtained from post translationally modified peptides while the other group had spectra obtained from peptides without any modifications.

**Figure 2:**
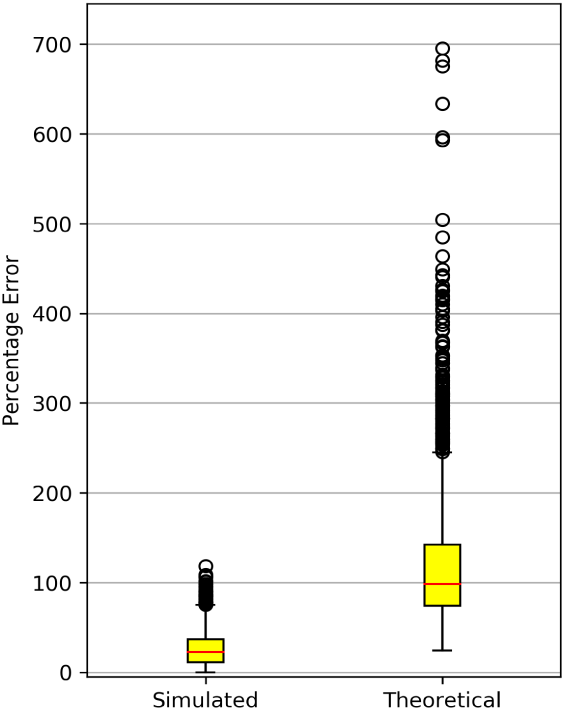
This box plot shows distribution of relative-error percentage for simulated spectra when compared against experimental spectra obtained from CID fragmentation and peptides without any post translational modifications

**Figure 3:**
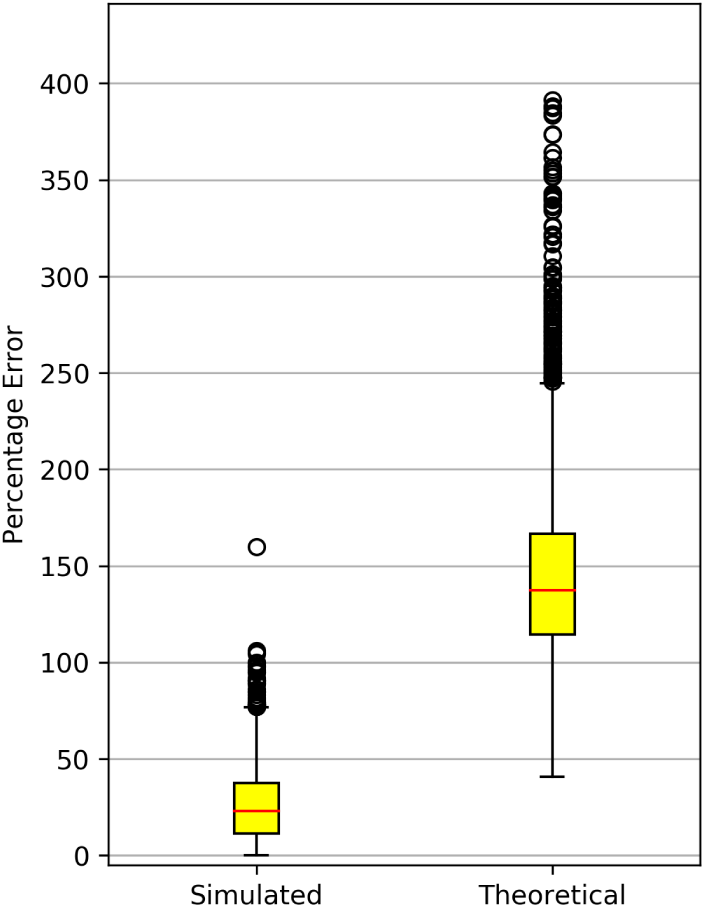
This box plot shows distribution of relative-error percentage for simulated spectra when compared against experimental spectra obtained from HCD fragmentation and peptides without any post translational modifications

**Figure 4:**
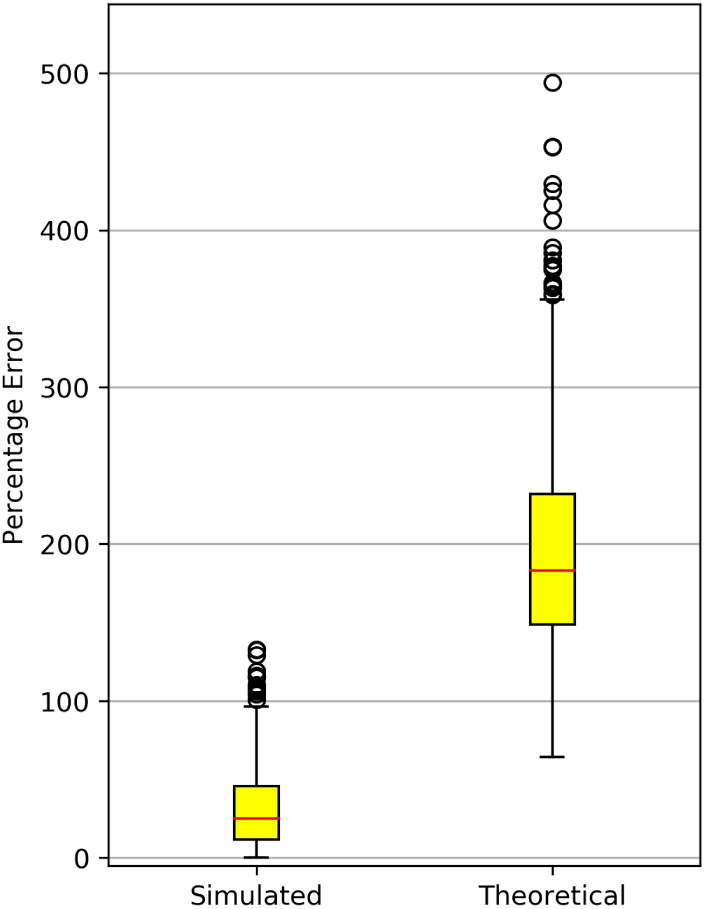
This box plot shows distribution of relative-error percentage for simulated spectra when compared against experimental spectra obtained from HCD fragmentation and peptides with post translational modifications

**Figure 5:**
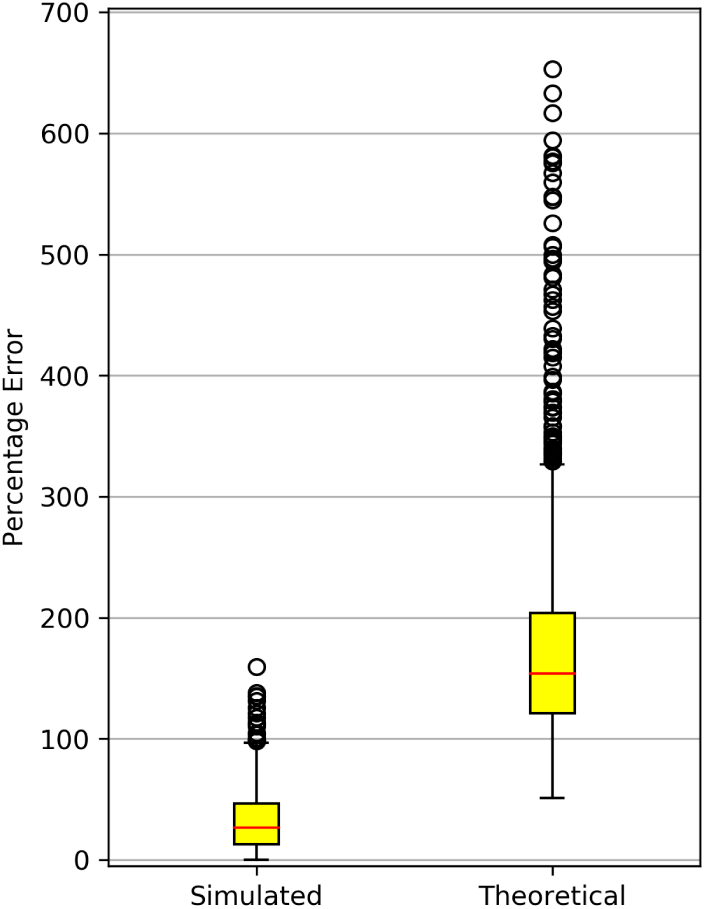
This box plot shows distribution of relative-error percentage for simulated spectra when compared against experimental spectra obtained from CID fragmentation and peptides with post translational modifications

It can be observed that the simulated spectra match the xcorr scores of experimental spectra much more closely than the theoretical spectra. Almost 75% simulated spectra have an error percentage of 25% which is extremely small compared to the large error percentage for theoretical spectra. Further, it can be observed that the error percentage remains consistent regardless of the type of ion-dissociation technique used or if the peptides were modified or not.

## 5 Assessment of Tide Using Simulated Spectra

In this section we demonstrate an application of MaSS Simulator by testing the performance of Crux Tide [16] peptide search engine by running it on 25 simulated MS/MS datasets with varying Peptide Coverage and SNR. The results show that varying SNR and Peptide Coverage affects the number of corrects hits by the database search software.

We obtained a yeast (*Pichia Pastoris*) proteome from uniprot.org (February 2018), it had a total of 5, 073 proteins in it. We performed an *insilico* complete tryptic digestion of the proteome with 3 missed cleavages allowed. This digestion was performed using the Protein Digestion Simulator [18]. Resulting peptides were filtered to shortlist peptides only with a mass between 800 and 4000 Dalton. This provided us with over 500, 000 peptides. To limit the size and time of experiments we picked 50, 000 peptides at random for our experiments. These peptides were then fed into the MaSS Simulator to simulate MS2 spectra with varying SNR and Coverage. We generated a total of 25 datasets with 50, 000 MS/MS spectra in each. Details of the configuration file used for generation of this data are given in Table. 1. Intensity values and ion types were determined based on the studies in [8] [14].

The simulated spectra were then searched against the yeast protein database using Crux-Tide software [16]. The results were then compared against the ground truth set provided by the MaSS Simulator. Results show that increasing the amount of noise puts the software off-track and decreases the amount of correct hits. Similarly spectra with poor peptide coverage also gave poor number of hits. These results can be observed in the Fig. 6.

**Figure 6:**
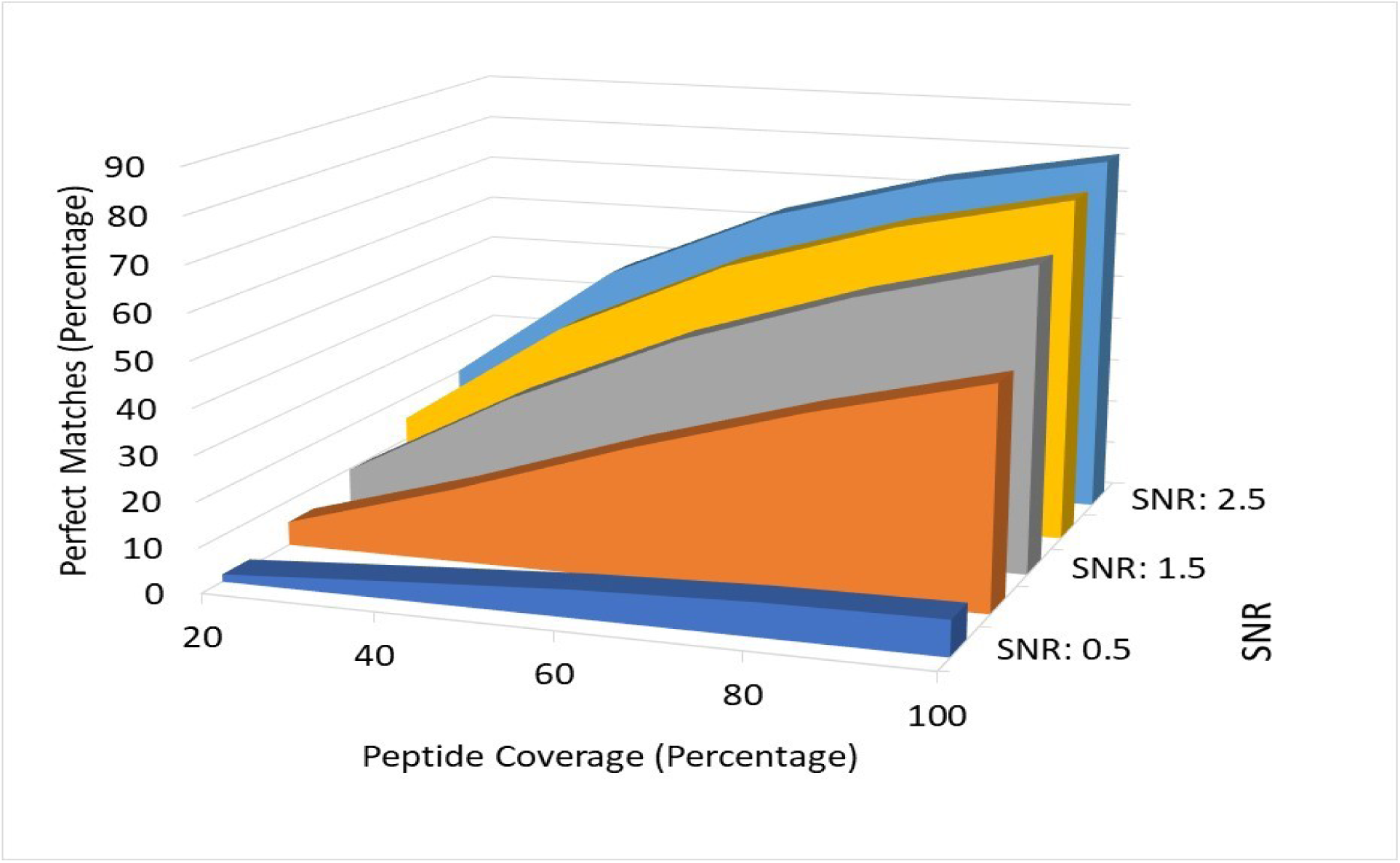
Figure showing assessment of Tide database search algorithm. It can be observed that the algorithm performs poorly on spectra with high noise and low coverage.

Using the simulated data and the ground peptides we were able to assess the accuracy of a given algorithm using a wide set of controlled conditions and parameters. This is not always possible when algorithms are only assessed using experimental MS data. We assert that the proposed simulator will be an indispensable tool for MS based proteomics method developers.

